# Inhibition of AXL dependent non-professional efferocytosis sensitizes pancreatic cancer to immune checkpoint blockade

**DOI:** 10.1101/2025.08.12.669943

**Authors:** Joon Park, Tyler McCaw, Hy Dao, Timothy Donahue, Joe Crompton, Erina Vlashi, Mark Girgis

## Abstract

AXL, a receptor tyrosine kinase, is upregulated in the majority of pancreatic adenocarcinoma (PDAC) and is associated with aggressive tumor behavior. Canonically, AXL is essential for efferocytosis, or the immunologically silent phagocytic clearance of apoptotic cells. While crucial to ensure self-tolerance, efferocytosis in cancer removes tumor antigens and dampens the antitumor response to create a pro-tumorigenic environment. Here, we show that PDAC cells co-opt the AXL pathway to act as non-professional efferocytes and clear chemotherapy induced apoptotic cells from their environment *in vitro* and *in vivo*. Inhibition of AXL with the selective inhibitor BGB324 inhibits efferocytosis and increases apoptotic cell burden, immunogenic cell death, and CD8 T cell infiltration in gemcitabine treated tumors *in vivo*. Single cell RNA expression analysis reveal that AXL inhibited gemcitabine treated tumors are enriched in the immunecheckpoint responsive TCF1^+^ CD8 T cells and respond to anti-PD1 therapy *in vivo*. We establish a novel ability of pancreatic cancer cells to engage in non-professional efferocytosis in response to chemotherapy. Inhibiting this response can be leveraged to sensitize this classically immune quiescent cancer to immunotherapy.

## Introduction

Pancreatic ductal adenocarcinoma (PDAC) remains one of the deadliest cancers with limited treatment effect resulting in nearly equivalent annual rates of incidence and mortality^1,2^. Advancements in novel therapeutic approaches are urgently needed.

AXL is a cell surface receptor tyrosine kinase that is selectively upregulated in over 70% of PDAC and is associated with aggressive oncogenic behavior including tumor growth, epithelial to mesenchymal transition, therapeutic resistance, immune suppression and metastasis^3-8^. Accumulating evidence shows that AXL expression is linked to T cell exclusion from the tumor microenvironment in PDAC and multiple tumor histologies, which likely contributes to increased aggressiveness of these cancers^7,9,10^.

Canonically, AXL is essential for efficient clearance of apoptotic cells (ACs) by phagocytic cells, a process known as efferocytosis (from Latin root *effere*, ‘to bury the dead ‘)^11^. This normal cellular process is crucial for the turnover of apoptotic cells to maintain tissue homeostasis, resolve inflammation, and promote self-tolerance^12,13^. Mechanistically, ACs present the ‘eat me ‘signal, phosphatidylserine (PS), on the cell surface. PS binds to a ‘bridging molecule ‘, Gas6, and the PS-Gas6 complex is recognized by the AXL receptor on efferocytes. The formation of AXL-Gas6-PS complex results in the engulfment of PS-positive ACs through a GTPase/Rac1-dependent cytoskeletal rearrangement^11^. Following successful AC clearance, efferocytes secrete anti-inflammatory cytokines and promote wound healing. Consequently, deficiencies or blockades in efferocytosis result in accumulation of ACs, their secondary necrosis, and release of immunogenic signals, such as damage-associated molecular patterns (DAMPs), which lead to inflammatory and autoimmune disorders^14^. Therefore, tight regulation of the efferocytosis pathway is critical for resolving inflammation, preventing breaches in self-tolerance, and promoting wound healing of normal tissues.

In cancer, which is often likened to a chronic non-healing wound, activation of the efferocytosis pathway promotes a pro-tumorigenic environment^15,16^. Moreover, commonly used cytotoxic cancer therapies, such as radiation therapy and chemotherapy, induce apoptotic cell death, likely amplifying efferocytic activity in the tumor and clearance of ACs. This can result in suppression of anti-tumor immunity, eventual immune escape, and treatment resistance^15,17,18^.

While efferocytosis is classically performed by innate immune cells like macrophages and dendritic cells, recent findings suggest that epithelial cells can also engage in efferocytosis, termed non-professional efferocytosis^19-21^. Whether epithelial-derived PDAC cells also utilize the AXL-Gas6-PS axis to participate in non-professional efferocytosis, remains unknown. Given AXL ‘s canonical role in efferocytosis and its pronounced overexpression in the majority of PDAC, we hypothesized that PDAC cells utilize the AXL-Gas6-PS axis to perform non-professional efferocytosis in response to cytotoxic therapy, ultimately promoting immune evasion and therapy resistance.

For the first time, we provide evidence that PDAC cells engage in efferocytosis to clear ACs following cytotoxic therapy, *in vitro* and *in vivo*. Using a well-characterized, selective AXL inhibitor we establish that PDAC cells engulf ACs in an AXL-dependent manner and are responsible for the removal of the majority of ACs, exceeding even the capacity of classic phagocytes. We also show that AXL inhibition allows for AC accumulation and priming of the host anti-tumor surveillance. This results in increased tumor T cell infiltration that can be leveraged to potentiate immune checkpoint blockade and improve tumor control. Together, these findings reveal non-classical efferocytosis as a novel mechanism employed by PDAC cells to escape immune surveillance. Targeting of this pathway has the potential to sensitize this classically immune quiescent cancer to immunotherapy, especially in combination with apoptosis-inducing cytotoxic therapy.

## METHODS

### Cell Culture

PANC-1, SUIT-2, MIAPACA-2, CFPAC-1, ASPC-1, and KPC were provided by Dr. Timothy Donahue (University of California, Los Angeles, CA). All cells were cultured in Dulbecco ‘s Modified Eagle Medium (Gibco) with 10% fetal bovine serum (Gibco) in 37°C in 5% CO2. Cells were routinely tested for *Mycoplasma* contamination with the Mycoplasma PCR Kit (Applied Biological Materials).

### Animal Studies

All animal studies were approved by Institutional Animal Care and Use Committee. Mice were purchased from The Jackson Laboratory. Six-to 10-week-old C57BL/6 mice were injected subcutaneously in the flank with 5 × 10^5^ KPC, KPC-Blue Fluorescent Protein (BFP) or KPC-Green Fluoresecent Protein (GFP) cells in 100µL PBS. Tumor volumes were measured using a caliper and estimated using the formula volume = (length x width^2^)/2. BGB324 (MedChemExpress), a selective AXL inhibitor, was resuspended in 0.5% hydroxypropyl methylcellulose/0.1% Tween80 solution and delivered via oral gavage twice daily at 50mg/kg for the treatment period. Gemcitabine (MedChemExpress) was resuspended in saline and delivered via intraperitoneal injection twice weekly at 50mg/kg for the treatment period. Mice were euthanized when the tumors reached 500mm^3^ or became ulcerated. Anti-PD1 antibody (BioXCell) or isotype control (BioXCell) was delivered via intraperitoneal injection at 10mg/kg twice weekly for the treatment period.

### Efferocytosis Studies

#### Induction of apoptosis

To induce apoptosis, cells in culture were placed in a 56°C incubator for 30 minutes. Cells were collected, pelleted, and washed twice with cold PBS, followed by labelling with pHrodo (Invitrogen) for 15 minutes, washed with cold 2% FBS in PBS and used for downstream applications.

#### Apoptosis/Necrosis Assay

The PI/Annexin kit (Invitrogen) was used per manufacturer guidelines and analyzed with flow cytometry to confirm apoptosis.

#### In vitro efferocytosis assays

To measure phosphatidylserine dependent uptake, untreated cells were labeled with CellTrace violet dye (Invitrogen) and co-cultured with GFP-phosphatidylserine (Echelon) or GFP-phosphatidylcholine beads (Echelon) in a 1:5 ratio for 6 hours. The cells were harvested, and uptake was measured with flow cytometry as percent GFP(+)/CellTrace(+). To measure apoptotic cell uptake, untreated cells were labeled with CFSE (Invitrogen) and co-cultured with pHrodo-stained apoptotic cells in a 1:5 ratio. The cells were harvested, and apoptosis was measured with flow cytometry as percent pHrodo(+)/CFSE (+).

#### Ex vivo efferocytosis assay

Single cell suspensions of digested mouse tumor were prepared (see “Tumor Digestion” below). The MACS dead cell removal kit (Miltenyi Biotec) was used to remove dead cells per manufacturer instructions. The cell suspension was plated in a 1:5 ratio with GFP-PS or GFP-PC beads for 6 hours. The cells were then harvested and prepared for flow cytometry staining and analysis.

#### In vivo efferocytosis assay

pHrodo-labeled apoptotic cells (50µL in PBS, 1 × 10^6^ cells) were delivered via intratumoral injection into subcutaneous syngeneic KPC tumors. After 6 hours, KPC tumors were harvested, and single cell suspensions were prepared and analyzed with flow cytometry.

### Immunohistochemistry

Tumors harvested from mice were formalin fixed and paraffin embedded. Formalin-fixed, paraffin-embedded tumor samples were incubated at 60 °C for 1 h, deparaffinized in xylene, and rehydrated with graded alcohol washes. Slides were then boiled in 0.01 M sodium citrate buffer for 15 min, followed by quenching of endogenous peroxidase with 3% hydrogen peroxide for antigen retrieval. After 1 h of blocking with 5% donkey serum at room temperature, primary antibodies were added and incubated overnight at 4 °C. Biotin-conjugated anti-rabbit secondary antibody (1:500; Jackson Labs) was added and developed using Elite Vectastain ABC kit.

### Incucyte

pHrodo labeled tumor cells were plated in 96 well plate in various drug concentrations and placed immediately in the Sartorious IncuCyte SX5 system. The IncuCyte system was used to obtain images at the specified timepoints and analyzed for red fluorescence using the IncuCyte software.

### Tumor Digestion

Tumors were digested using tumor digestion buffer consisting of 2 mg/mL collagenase Type IV (Gibco), 0.2mg/mL DNAse I (Invitrogen), and 1mg/mL Dipase II (Gibco) in Roswell Park Memorial Institude Medium (Gibco). Tumors were minced into less than 2mm chunks and digested at 37°C for 30 minutes, filtered through a 70µM strainer, pelleted, washed and resuspended in single cell suspension for downstream analysis.

### Lentivirus Transduction

GFP and BFP lentivirus vectors were obtained from UCLA Molecular Screening Shared Resource. KPC cells were infected with the lentivirus in 10% DMEM with 8µg/mL polybrene overnight. Flow cytometry sort on the transduced cells was used to select the top 1% of stably transduced cells.

### Flow Cytometry

Single cell suspensions were prepared as above and stained with Zombie Viability dye (BioLegend) for 20 minutes in room temperature (RT) in the dark. The cells were then washed with cold 1:250 EDTA, 2% FBS in PBS and incubated with Fc Block (BioLegend) and extracellular antibodies for 30 minutes. For intracellular staining, cells were fixed and permeabilized using theCytofix/Cytoperm kit (BD Biosciences) per manufacturer instructions. Cells were stained with an intracellular antibody cocktail for 1 hour in the dark, washed, and analyzed with the Attune NxT Flow Cytometer or ImageStreamx MarkII Imaging Flow Cytometer. OMIQ (Dotmatics) was used for data analysis.

### Single Cell RNA Library Preparation and Sequencing

Tumors were digested and prepared in single cell suspension solution as above. Cells were stained with Zombie live/dead and CD45 and sorted using flow sorter for Live CD45+ (BioLegend) cells. These cells were suspended in PBS for single-cell library creation. Samples were counted using Countess II Automated Cell Counter (Thermo Fisher Scientific) and hemocytometer for cell concentration using Trypan Blue stain 0.4% (Invitrogen). Single cell gene expression libraries were created using Chromium Next GEM Single Cell 3 ‘(v3.1 Chemistry) (10x Genomics), Chromium Next GEM Chip G Single Cell Kit (10x Genomics), and Dual Index Kit TT, Set A (PN-1000215) (10x Genomics) according to the manufacturer ‘s instructions. Briefly, cells were loaded to target 5,000 cells to form GEMs and barcode individual cells. GEMs were then cleaned. cDNA and libraries were also created according to manufacturer ‘s instructions. Library quality was assessed using 4200 TapeStation System and D1000 ScreenTape (both from Agilent) and Qubit 2.0 (Invitrogen) for concentration and size distribution. Samples were sequenced using Novaseq X Plus sequencer (Illumina) using 100 cycles. 150M reads were targeted for each sample, targeting 40,000 reads per cell.

### Single Cell RNA Data Analysis

The 10x Genomics Cell Ranger 7.1.0 program was used to align the raw data to the mm10-2020-A mouse reference transcriptome. The R package Seurat was used for data preprocessing and analysis. Low quality reads were filtered out using cutoffs of percent.mt <20, nfeature <200 and ncount<500. The log transform pipeline in Seurat was used to normalize the data and clusters were identified using the FindClusters function using a resolution of 1.2. The CellDex package was used to call the Immunologic Genome Project data and the SingleR package was used to annotate the clusters. The clusters were then further manually annotated using known marker genes. UMAPs were generated using the Seurat package.

### Statistical Analysis

Data are presented as means ±SD with indicated biologic replicates. Comparisons of two groups were calculated using unpaired two-tailed Student *t* test, and *p* values less than 0.05 were considered significant. All analyses were conducted by using Prism version 10.4.1 or R-Studio version 4.3.1. Pairwise tumor growth curves comparisons were performed using mixed-effect modeling with log transformation using the *TumGrowth* package in R-Studio. ^22^

## Data Availability

All data are available on request from the corresponding author

## Results

### Pancreatic cancer cells efferocytose apoptotic cells in an AXL dependent fashion

Live human PDAC cells were co-cultured with apoptotic PDAC cells (ACs) to determine whether the live cells engulf ACs via efferocytosis. Heat-induced ACs were labeled with the pH-sensitive, red-fluorescing dye pHrodo, which only fluoresces under acidic conditions, such as after engulfment into a phagolysosome. Apoptosis was confirmed via the co-stain with propidium iodide (PI) and Annexin V (Supp. Fig. 1A). pHrodo-labeled ACs were then co-cultured with live PDAC cells labeled with the CellTrace green-fluorescing dye, carboxyfluorescein diacetate succinimidyl ester (CellTrace CFSE) (Fig. 1A). Single cell imaging flow cytometry (ImageStream) analysis showed that CFSE-labeled, live PDAC cells readily engulf pHrodo-labeled ACs, clearly visualized as red-fluorescent bodies inside acidic intracellular vesicles in the cytoplasm of live PDAC cells (Fig. 1B). Pre-incubation with an efferocytosis inhibitor, Zinc69391, or phosphatidylserine (PS) masking antibody, Annexin V, significantly reduces the fraction of CellTrace^+^/pHrodo^+^ cells in the co-culture (Fig. 1C-D, Supp. Fig. 1B), demonstrating effective inhitibion of efferocytosis^23^. The efferocytic acitivity is depedent on AXL, as pre-incubation with the AXL inhibitor, BGB324 dose-dependently reduced the fraction of CellTrace^+^/pHrodo^+^ efferocytic cells (Fig. 1E, blue vs. green bars, Supp. Fig. 1C) in six different human and one murine PDAC line. This cell panel also revealed that ∼15-30% of PDAC cells exhibit efferocytic acitivity *in vitro* at baseline (Fig. 1E, blue bars).

**Figure 1.**
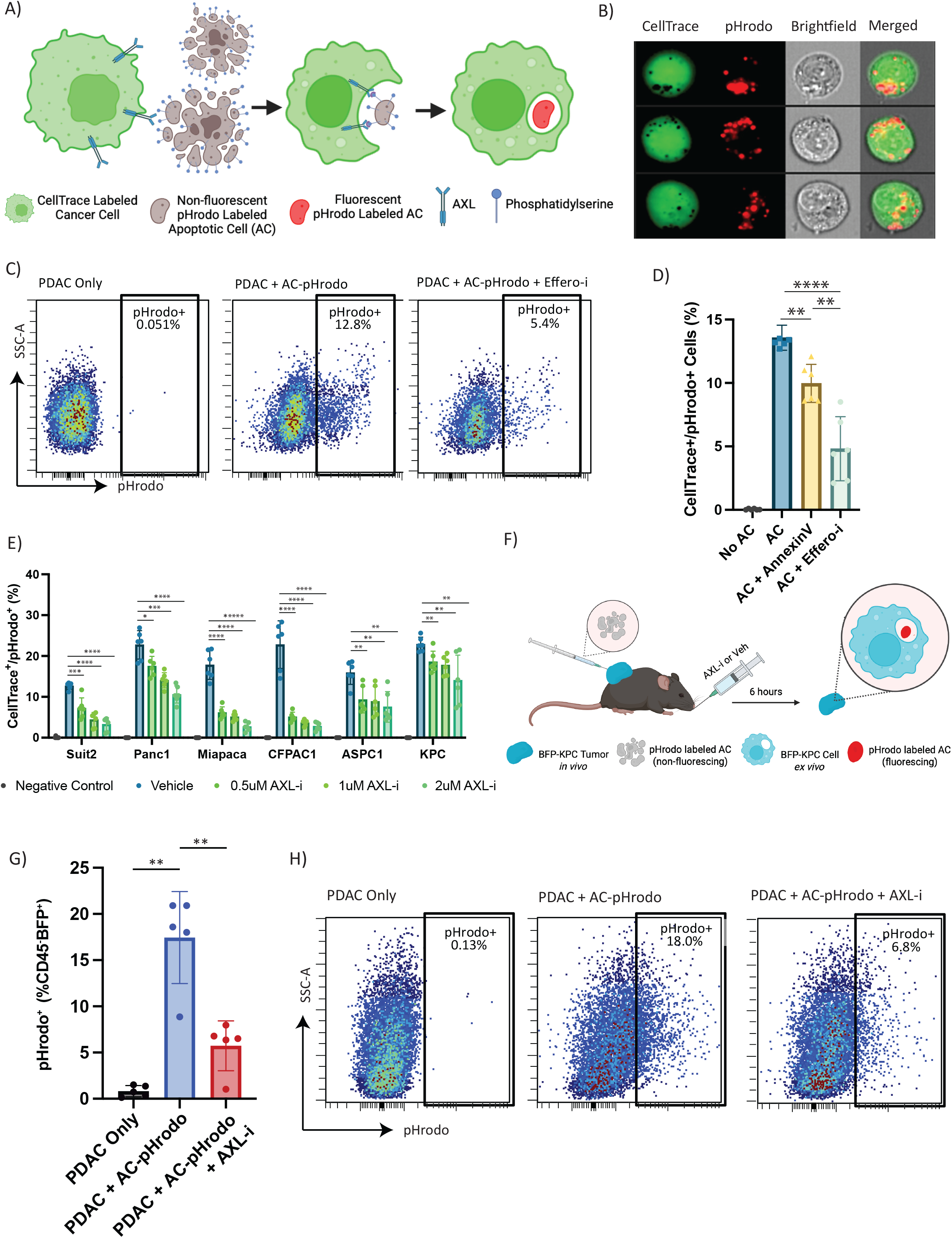
Pancreatic cancer cells efferocytosis apoptotic cells in an AXL dependent fashion. (A) SUIT2 PDAC cells were labeled with CFSE and co-cultured with pHrodo labeled apoptotic cells for 6 hours. (B) Images representative of the CFSE+ pHrodo+ population visualized on ImageStream Flow Cytometry. (C,D) Percent pHrodo+ of CFSE+ cells after treatment with phosphatidylserine antibody Annexin V, efferocytosis inhibitor Zinc69391, or control (n=5-6). ** p<0.01. ****p<0.0001. (E) Effect of increasing doses of AXL inhibitor BGB324 on engulfment activity across a panel of pancreatic cell lines (n=6). (F) pHrodo labeled apoptotic cells were injected into inoculated BFP expressing KPC tumors in C57BL/6 mice pretreated with 50mg/kg BGB324 or vehicle and measured for pHrodo activity after 6 hours. (G) The effect of AXL inhibitor BGB324 on pHrodo apoptotic cell uptake by BFP-KPC cells *in vivo* (n=5). **p<0.01. (H) Flow plots representative of pHrodo fluorescence in the CD45-BFP+ cell population.

To demonstrate efferocytic activity *in vivo*, we subcutaneously implanted the murine KPC cells stably expressing blue fluorescent protein (BFP) in C57BL/6J mice. After tumors were established, mice were treated with the AXL inhibitor or vehicle and pHrodo-labeled apoptotic KPC cells were injected intratumorally 12 hours after treatment (Fig. 1F). Six hours following apoptotic cell injection, the tumors were removed, processed into single cell suspensions and stained with CD45 to exclude white blood cells from the flow cytometry analysis. Similar to *in vitro* findings, ∼15-20% of CD45^-^BFP^+^ KPC tumor cells co-stain with pHrodo, demonstrating that approximately a quarter of the KPC cells *in vivo* actively engulf pHrodo^+^ ACs in an AXL-dependent manner (Fig. 1G-H). Together, these results support the conclusion that live PDAC cells can perform efferocytosis *in vitro* and *in vivo* to engulf ACs in an AXL-dependent fashion.

### Pancreatic cancer cells perform efferocytosis in a phosphatidylserine-specific manner

Efferocytosis is a highly regulated process involving the recognition of the apoptotic ‘eat me ‘phosphatidylserine (PS) signal on the cell surface. To demonstrate the specific involvement of PS in PDAC efferocytosis, unlabeled human PDAC cells were co-cultured with green fluorescent protein (GFP)-labeled PS or sham control GFP-phosphatidylcholine (PC) beads (Fig. 2A). In line with PS-specific efferocytosis, PDAC cells selectively engulfed PS beads, but not PC beads, in an AXL-dependent fashion (Fig. 2B), at frequencies (∼20%) similar to those observed in AC co-cultures (Fig. 1).

**Figure 2.**
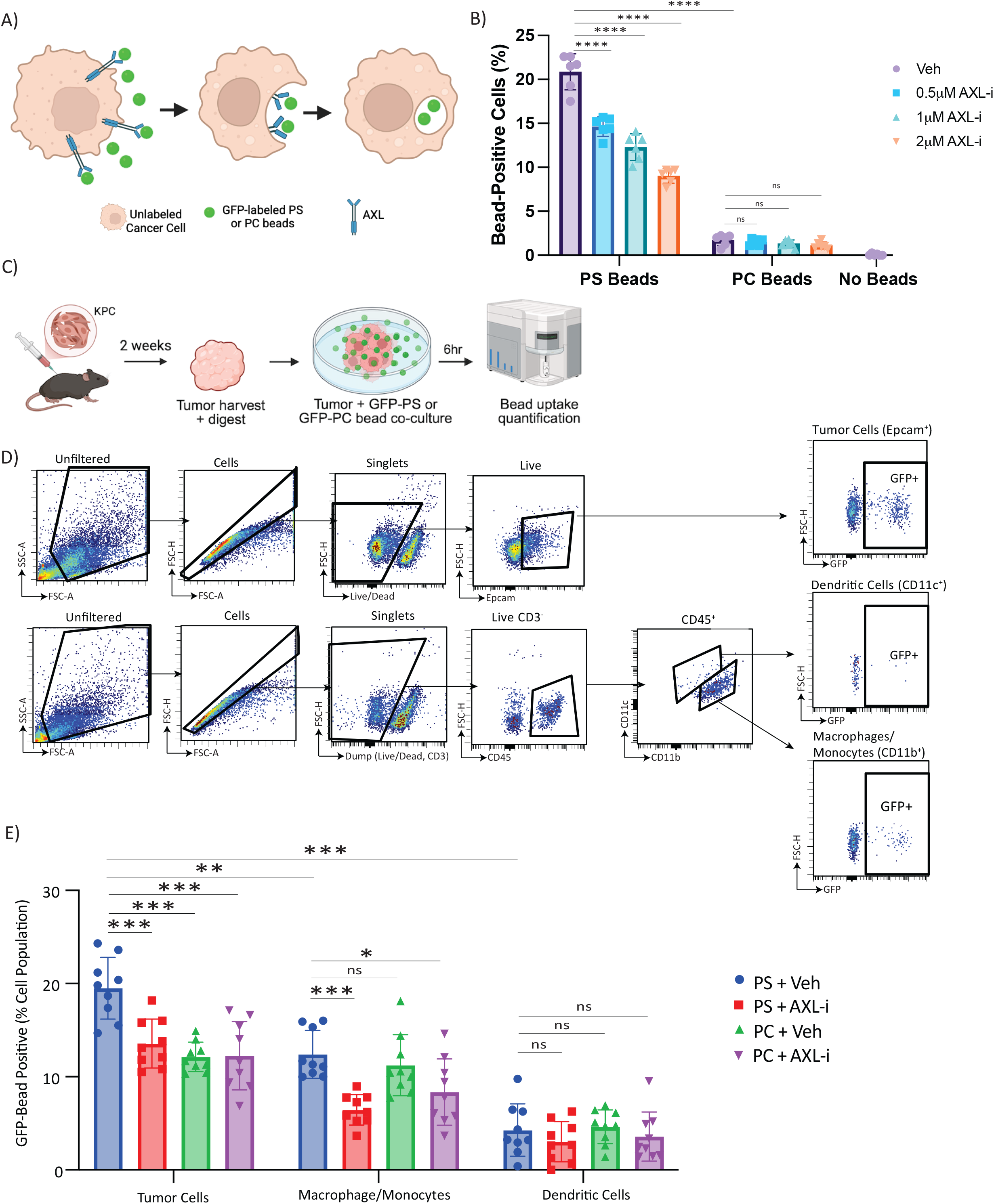
Pancreatic cancer cells perform efferocytosis in a phosphatidylserine-specific manner. (A,B) Phosphatidylserine GFP particle or control phosphatidylchloline GFP particle uptake in 6 hour coculture with unlabeled SUIT2 cells treated with increasing doses of AXL inhibitor BGB324 (n=6). ****p<0.0001. (C) Inoculated KPC tumors from C57BL/6 mice dissociated into single cell suspension and co-cultured with PS-GFP or PC-GFP particles with AXL inhibitor BGB324 or vehicle. (D) Flow gating strategy for tumor cells, dendritic cells, and macrophages. (F) Effect of AXL-inhibitor on PS-GFP or PC-GFP particle uptake by tumor cells, macrophages and dendritic cells (n=9). ***p<0.001. *p<0.05.

The KPC murine PDAC model provides the opportunity to compare the efferocytic activity of pancreatic cancer cells relative to professional phagocytes, such as macrophages or dendritic cells *in vivo*. For this purpose, unlabeled KPC subcutaneous tumors were digested into single cell suspensions and co-cultured with GFP-labeled PS or PC beads *ex vivo* (Fig. 2C). Flow cytometry was used to immunophenotype cellular subpopulations in the tumor microenvironment. Tumor cells were defined as EPCAM^+^, dendritic cells were defined as CD45^+^CD11c^+^CD11b^-^ and monocytes/macrophages were defined as CD45^+^CD11b^+^CD11c^-^ (Fig. 2D). The *ex-vivo* co-culture showed a similar pattern of bead uptake to the *in vitro* co-culture (Fig. 2A-B) with ∼20% of tumor cells selectively engulfing PS beads in an AXL-dependent fashion (Fig. 2E). There was some nonspecific PC bead uptake by the tumor cells that was unaffected by AXL inhibition (Fig. 2E). In contrast, macrophages/monocytes and dendritic cells equally took up PS and PC beads, consistent with their nonspecific phagocytic function.

Notably, tumor cells showed a significantly higher percentage of PS bead uptake relative to macrophages/monocytes (*p*<0.001) or dendritic cells (*p<*0.0001) (Fig. 2E).

### Gemcitabine increases efferocytic activity of pancreatic cancer cells

Most cytotoxic therapies used to treat cancers induce apoptotic cell death. Therefore, we investigated whether the chemotherapeutic agent gemcitabine, which is commonly used clinically to treat patients with PDAC, increases efferocytic activity by PDAC cells. To simplify the labeling for detection of efferocytic activity, the human SUIT2 and the murine KPC cell lines were stained with pHrodo and exposed to increasing doses of gemcitabine and 2µM of AXL inhibitor, *in vitro*. Since pHrodo only fluoresces in acidic conditions such as in the phagolysosome, pHrodo-positive cells represent live cells that have engulfed gemcitabine-induced apoptotic cells (Fig. 3A). The percent of pHrodo^+^ cells increases in a gemcitabine dose-dependent manner, suggestive of increased efferocytic activity (Fig. 3B-C, Supp. Fig. 2B), in an AXL-dependent manner, in keeping with the efferocytosis of heat-induced ACs.

**Figure 3.**
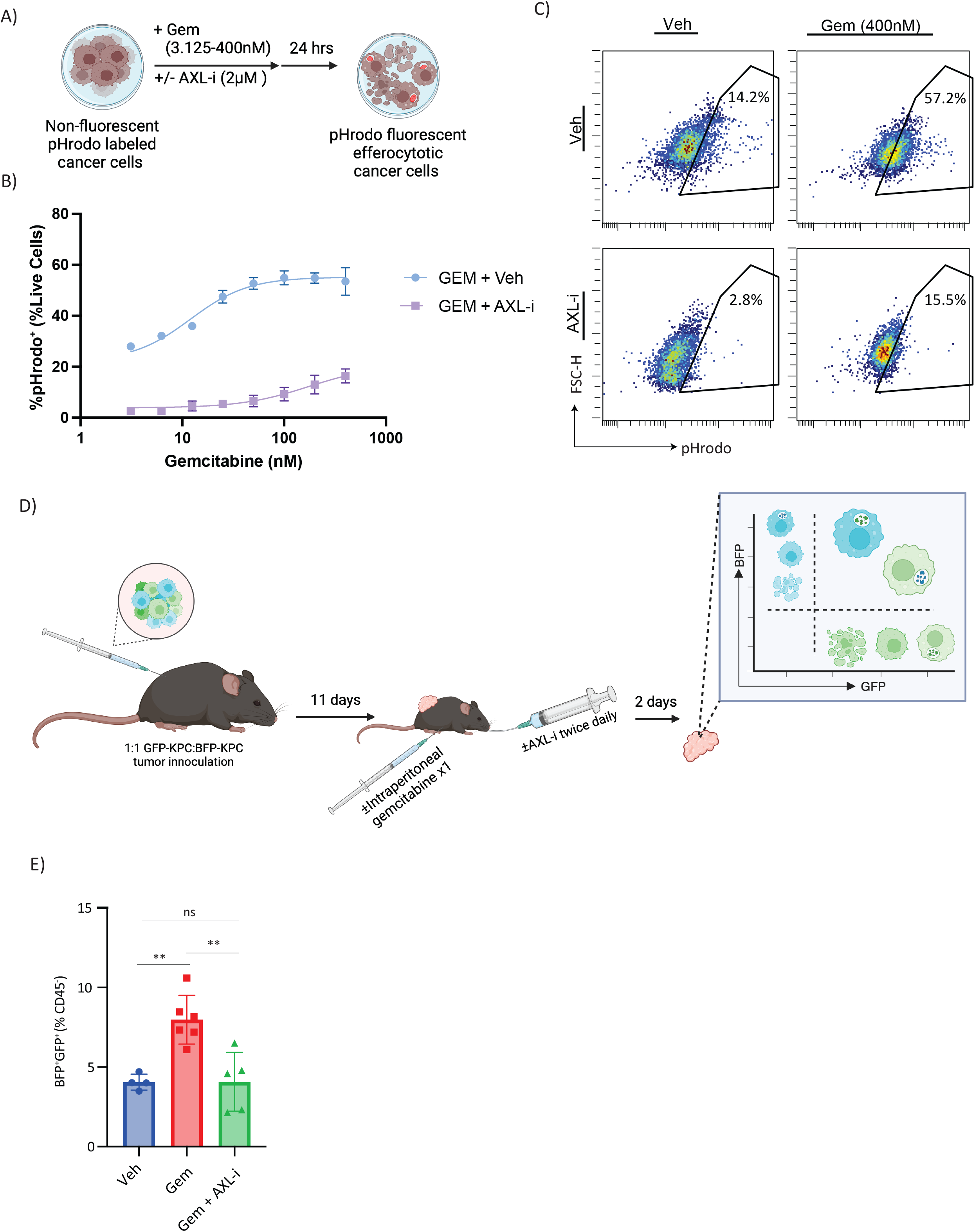
Gemcitabine increases efferocytic activity of pancreatic cancer cells. (A) Schematic. pHrodo labeled KPC cells were treated with gemcitabine doses ranging from 3.125nM-400nM with or without 2µM BGB324 and measured for fluorescence with flow cytometry. (B) Percent pHrodo^+^ KPC cells with increasing gemcitabine doses with or without AXL-i BGB324 and (C) representative flow plots. (D) Flow analysis was performed on single cell suspensions prepared from C57BL/6 mice inoculated with a 1:1 mixture of GFP-KPC:BFP-KPC tumors treated with 50mg/kg gemcitabine intraperitoneally with or without 50mg/kg AXL-i BGB324 via oral gavage. (E) The effect of gemcitabine on tumor cell efferocytosis activity compared to vehicle and with AXL inhibition (n=4-6). **p<0.01. ns=not significant.

Since pHrodo fluorescence is diluted to background levels after a few cell divisions and therefore not suitable for tracking efferocytosis in an *in vivo* model, we used stably transduced BFP-KPC and GFP-KPC cells to characterize the effect of gemcitabine on AXL-dependent efferocytosis *in vivo*. C57BL/6 mice were implanted with a 1:1 ratio of GFP- and BFP-labeled KPC tumor cells. After tumors were established, mice were administered one dose of gemcitabine at 50mg/kg and AXL inhibitor at 50mg/kg, twice daily for 48 hours. Single cell suspensions of tumors were used to characterize the fraction of GFP^+^BFP^+^ efferocytic cells within the CD45^-^ population via flow cytometry (Supp. Fig 2C). CD45^+^ cells were excluded to eliminate any professional phagocytes that had engulfed BFP^+^ and GFP^+^ apoptotic tumor cells. Gemcitabine treatment increased the efferocytic BFP^+^GFP^+^ population in the tumor by 2-fold and AXL inhibition prevented this increase (Fig. 3D-E).

### AXL inhibition promotes an immunogenic cell death response to cytotoxic therapy

Activation of the efferocytosis pathway leads to an immunosuppressive environment in cancer due to antigen removal and inhibition of T cell responses^15^. Conversely, inhibition of efferocytosis results in accumulation of apoptotic cells, release of DAMPs following secondary necrosis, and increased immunogenicity. Given our findings that PDAC cells contribute significantly to clearance of apoptotic cells in *in vitro* and *in vivo*, at least in part via the AXL-PS axis (Fig. 1-3), we sought to understand the effect of AXL inhibition on the PDAC immune landscape, with or without gemcitabine induced cell death. KPC tumor-bearing C57BL/6 mice were treated with gemcitabine twice weekly and AXL inhibitor twice daily (Fig. 4A). As expected, treatment with gemcitabine reduced tumor growth, AXL inhibition alone had no impact on established tumors, and AXL inhibition had no additional benefit on tumor control after 1 week of combined treatment (Fig. 4B,C). However, tumors treated with the combination approach had significantly higher fractions of apoptotic cells as measured immunohistochemically by cleaved caspase 3 (Fig. 4D). Notably, this increased fraction of apoptotic cells was accompanied by an increase in DAMPs as measured by HMGB1 and a corresponding increase in CD8 T cell infiltration (Fig. 4D), suggestive of increased engagement in antitumor immunity. Consistent with these results, flow cytometry performed with similarly treated tumors confirmed a significantly increased CD8+ population following AXL-i monotherapy and combination therapy (Fig. 4E).

**Figure 4.**
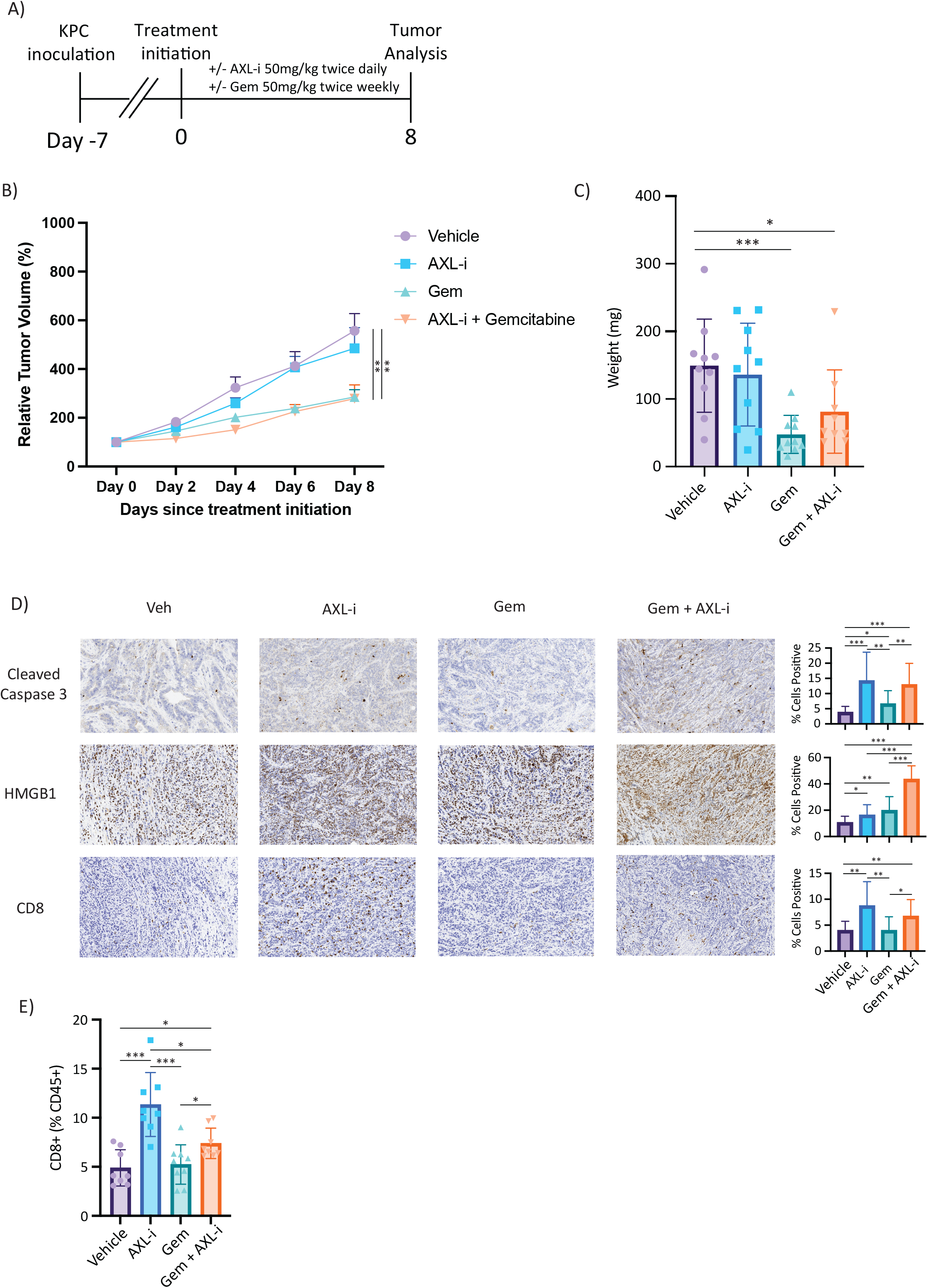
AXL inhibition promotes an immunogenic cell death response to cytotoxic therapy. (A) C57BL/6 mice inoculated with KPC cells were treated with AXL-i BGB324 twice daily and gemcitabine twice weekly for 8 days and measured for (B) tumor volume and (C) tumor weight (n=9-10). *p<.05. **p<0.01. ***p<0.001. (D) Immunohistologic staining of cleaved caspase 3, HMGB1, and CD8 in KPC tumors from mice treated with gemcitabine and AXL-i BGB324 (n=4). *p<.05. **p<0.01. ***p<0.001. (E) Flow cytometry of KPC tumors in C57BL/6 mice treated with gemcitabine ± AXL-i BGB324 (n=8-9).

Single cell RNA sequencing analysis was performed on the flow sorted CD45^+^ population of the treated tumors to better characterize the immune reponse to combination therapy. Unsupervised clustering revealed three distinct CD8^+^ populations: progenitor exhausted, proliferating and terminally exhausted (Fig. 5A)^24^. The stem-like T progenitor exhausted population was defined by the high *Tcf7* and *Il7r* expression and the T terminally exhausted population was defined by high expression of the inhibitory markers *Pdcd1, Ctla4, Lag3, Havcr2*, and *Entpd1*. The T proliferating population had moderate expression of inhibitory markers and high expression of *Mki67* (Fig. 5B). Single cell analysis again showed increased CD8 populations with AXL-i monotherapy and combination therapy (Fig. 5C,D). Subpopulation analysis revealed that AXL-i monotherapy increased CD8^+^ population mainly by inducing an increase in terminally exhausted subpopulation of T cells. In contrast, treatment with AXL-i combined with gemcitabine expanded the subpopulation of stem-cell like T progenitor population (Fig. 5C,D).

**Figure 5.**
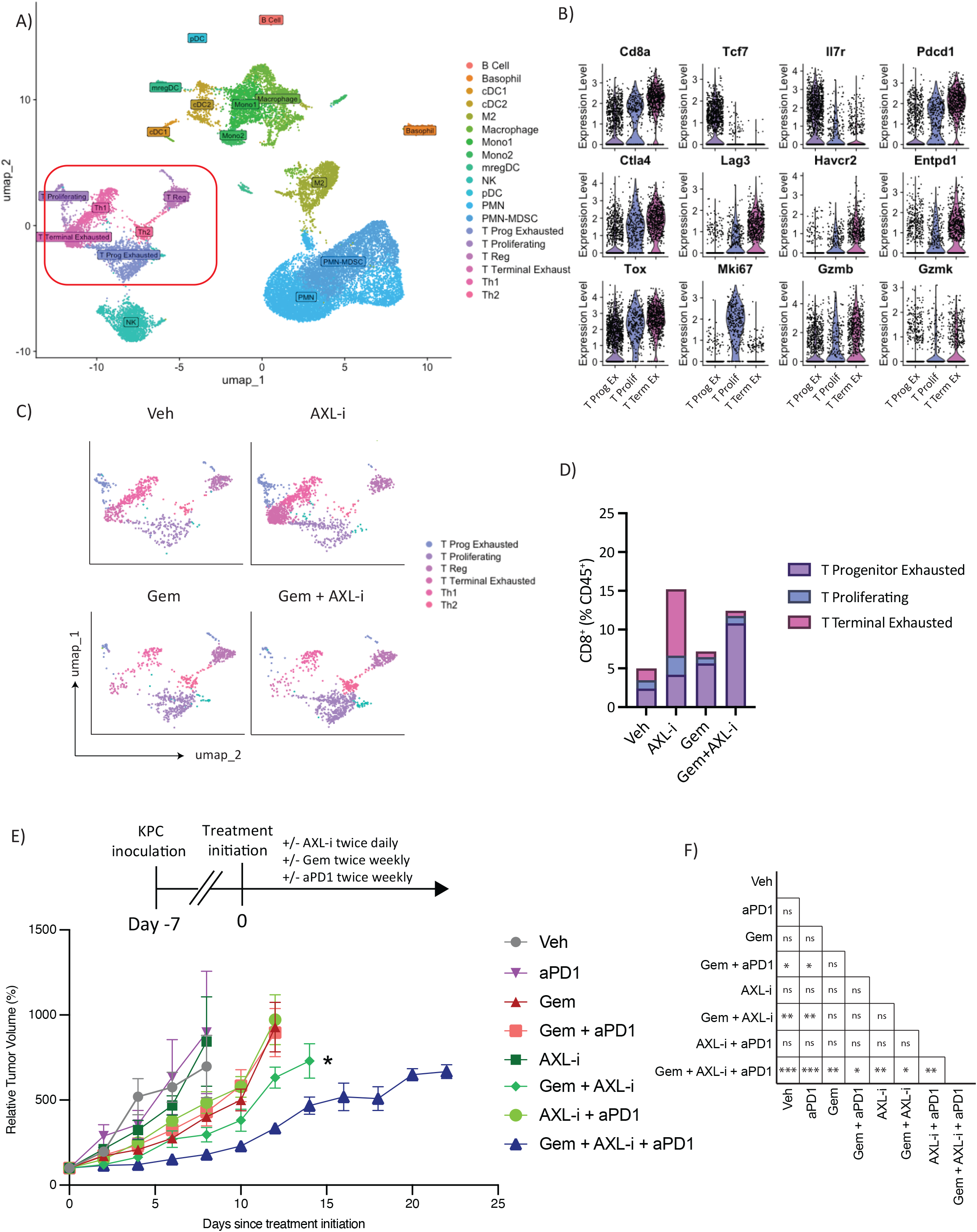
Combining AXL inhibition with gemcitabine expands the T progenitor exhausted population and sensitizes pancreatic cancers to anti-PD1 therapy. (A) UMAP plot of pooled 29875 sequenced CD45+ sorted cells in 19 clusters from C57BL/6 mice treated with vehicle, AXL-i BGB324 50mg/kg twice daily, gemcitabine 50mg/kg twice weekly or combination therapy for one week (n=2/group). The CD3 population is highlighted. (B) Relative expression of marker genes used to annotate CD8 subset populations. (C) UMAP plot and (D) bar plot showing the distribution of CD3 cells by treatment group. (E,F) Survival curves and pairwise comparisons of C57BL/6 mice inoculated with KPC cells treated with combinations of gemcitabine 50mg/kg twice intraperitoneally weekly, AXL-i BGB324 50mg/kg oral gavage twice daily, and anti-PD1 antibody 10mg/kg intraperitoneally twice weekly. ns=not significant. *p<.05. **p<0.01. ***p<0.001.

### Combining AXL inhibition with gemcitabine sensitizes pancreatic cancers to anti-PD1 therapy

Studies have shown that exhausted progenitor T cells, characterized transcriptionally as *Tcf7*^+^ CD8, are responsive to anti-PD1 therapy^25-27^. Our findings of an expanded *TCF7*^*+*^ CD8 population in PDAC tumors treated with AXL-i plus gemcitabine led us to hypothesize that anti-PD1 therapy will further improve tumor control when combined with AXL inhibition and gemcitabine. In line with this, only tumors treated with both AXL inhibition and gemcitabine received significant benefit from anti-PD1 therapy (Fig. 5 E,F).

## Discussion

This study establishes a previously undescribed phenomenon of intrinsic efferocytic activity in pancreatic cancer cells and its role in modulating immunogenic cell death. We show that pancreatic cancer cells co-opt the AXL pathway, which is upregulated in the majority of PDAC, to clear apoptotic cells from their environment. This efferocytic activity is upregulated in settings of increased apoptotic burden, such as during chemotherapy. We also show that combining AXL inhibtion with chemotherapy *in vivo* is associated with an accumulation of ACs and release of DAMPs, expansion of progenitor exhausted T cells, and sensitization to anti-PD1 therapy.

Canonical AXL mediated efferocytosis clears apoptotic cells via AXL expressing phagocytes that recognize the ‘eat me ‘phosphatidylserine signal on apoptotic cells, facilitating tolerogenic cell death. Our results show that pancreatic cancer cells readily engulf apoptotic cells in their environment in an AXL dependent fashion *in vitro* and *in vivo*. The finding that this engulfment was phosphatidylserine specific is supportive of a regulated apoptotic cell clearance process, distinct from the well characterized phenomenon of macropinocytosis, in which pancreatic cancer cells nonspecifically engulf extracellular content primarily for nutrient scavenging^28,29^. Notably, a recent study on macropinocytosis by King et al. noted that AXL is required for the uptake of cell debris^30^.

We find that the increase in gemcitabine-induced apoptotic cell burden is associated with increased AXL-mediated efferocytic activity and exacerbated immunosuppressive microenvironment in pancreatic tumors, as measured by DAMPs and T cell exclusion. This is consistent with reports that gemcitabine treatment induces an immunosuppressive microenvironment^31,32^, likely in part due to the induction of tolerogenic cell death facilitated by the pancreatic cancer cells engaging in intrinsic efferocytic activity. Moreover, our *ex vivo* studies indicate that pancreatic cancer cells outperform macrophages and dendritic cells in uptaking PS-labeled beads, suggesting that tumor cells are the dominant efferocytes in the PDAC microenvironment. *In vivo* inhibition of AXL during gemcitabine treatment leads to an accumulation of unprocessed apoptotic cells in the microenvironment that favors an immunogenic cell death response due to secondary necrosis. This is evidenced by the increase in DAMP signaling in tumors treated with a combination of gemcitabine and AXL inhibitor, consistent with previous reports.^7,10^ Altogether, our findings support the thesis that the intrinsic, AXL-mediated efferocytosis activity of PDAC cells promotes an immunosuppressive microenvironment, which is exacerbated by apoptosis-inducing cytotoxic chemotherapeutics, such as gemcitabine. AXL blockade therefore, induces an “efferocytosis blockade shunt”^33^, forcing uncleared apoptotic cells to undergo secondary necrosis and release DAMPs and tumor antigen, which reveals them to the adaptive immune system, ultimately stimulating an antitumor immune response (Fig. 6).

**Figure 6.**
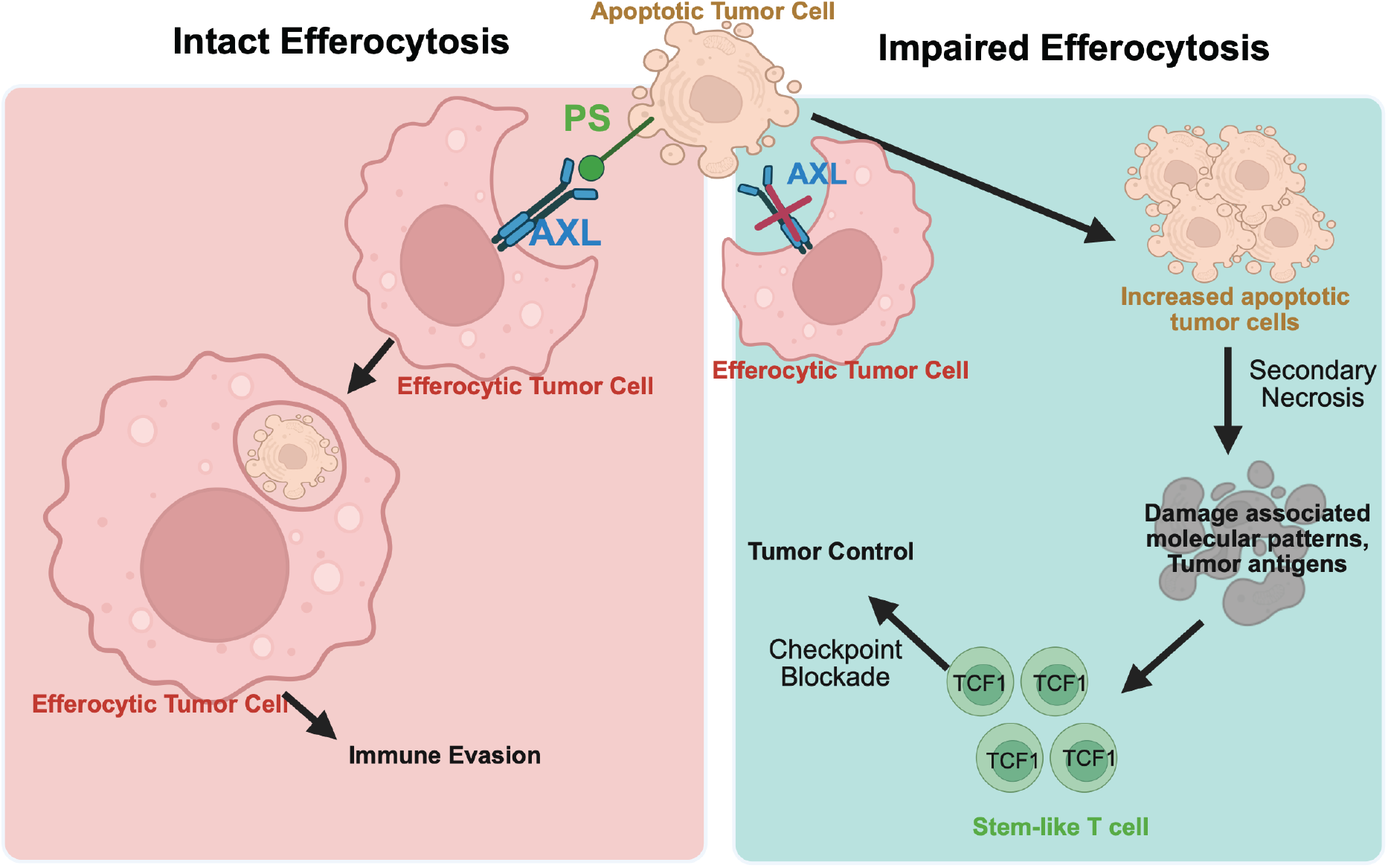
Visual Abstract. Pancreatic tumor cells engage in efferocytosis to hide apoptotic tumor cells from immune surveillance via AXL. Inhibition of AXL leads to increased apoptotic cell burden, their secondary necrosis and release of DAMPs, increased CD8 cell engagement and sensitization to anti-PD1 therapy.

Combining AXL inhibition with gemcitabine sensitizes PDAC to anti-PD1 therapy *in vivo*, while neither approach alone benefited from anti-PD1 therapy. This suggests that both, the increased input of apoptotic cells with cytotoxic therapy and decreased processing capacity by AXL inhibition is required to overwhelm the tumor ‘s ability to maintain innate tolerogenic cell death and shift towards adaptive immunogenic cell death. Mechanistically, anti-PD1 sensitization is explained by the expansion of the TCF1^+^ “progenitor exhausted” CD8 T cell population *in vivo* following combination therapy. This T cell subset serves as a long-lived stem-like reservoir that accounts for the proliferative burst with anti-PD1 therapy, asymmetrictrically dividing to produce effector-like T cells with cytotoxic activity. Our findings are consistent with a recent report that AXL inhibition expands the TCF1^+^PDL1^+^ cells in treatment resistant lung cancer *in vivo*, with a mechanistic rationale primarily centered around AXL expressing dendritic cells^34^. Given that AXL is expressed across many cell types in the tumor microenvironment with pleotropic activity, our proposed efferocytosis mechanism supplements without contradicting this prior report. To our knowledge, our study is the first to report pharmacologic induction of TCF1^+^ CD8 T cell expansion in PDAC, a much more classically immune quiescent tumor than even treatment resistant lung cancers^35^.

Our work describes a novel concept of tumor cell intrinsic efferocytosis activity that can be leveraged to shunt cytotoxic therapy induced apoptotic cells toward immunogenic cell death and sensitize pancreatic cancers to immune directed therapy. While our study provides a novel mechanism supporting the growing number of reports that AXL alters the immune landscape in PDAC, further studies are required to quantify the relative contributions of different AXL expressing cells in expanding the TCF1^+^ CD8 population, as well as address whether other commonly used cytotoxic therapies in PDAC such as FOLFIRONIX and radiation have similar effects.

---

**Supplementary Figure 1.**
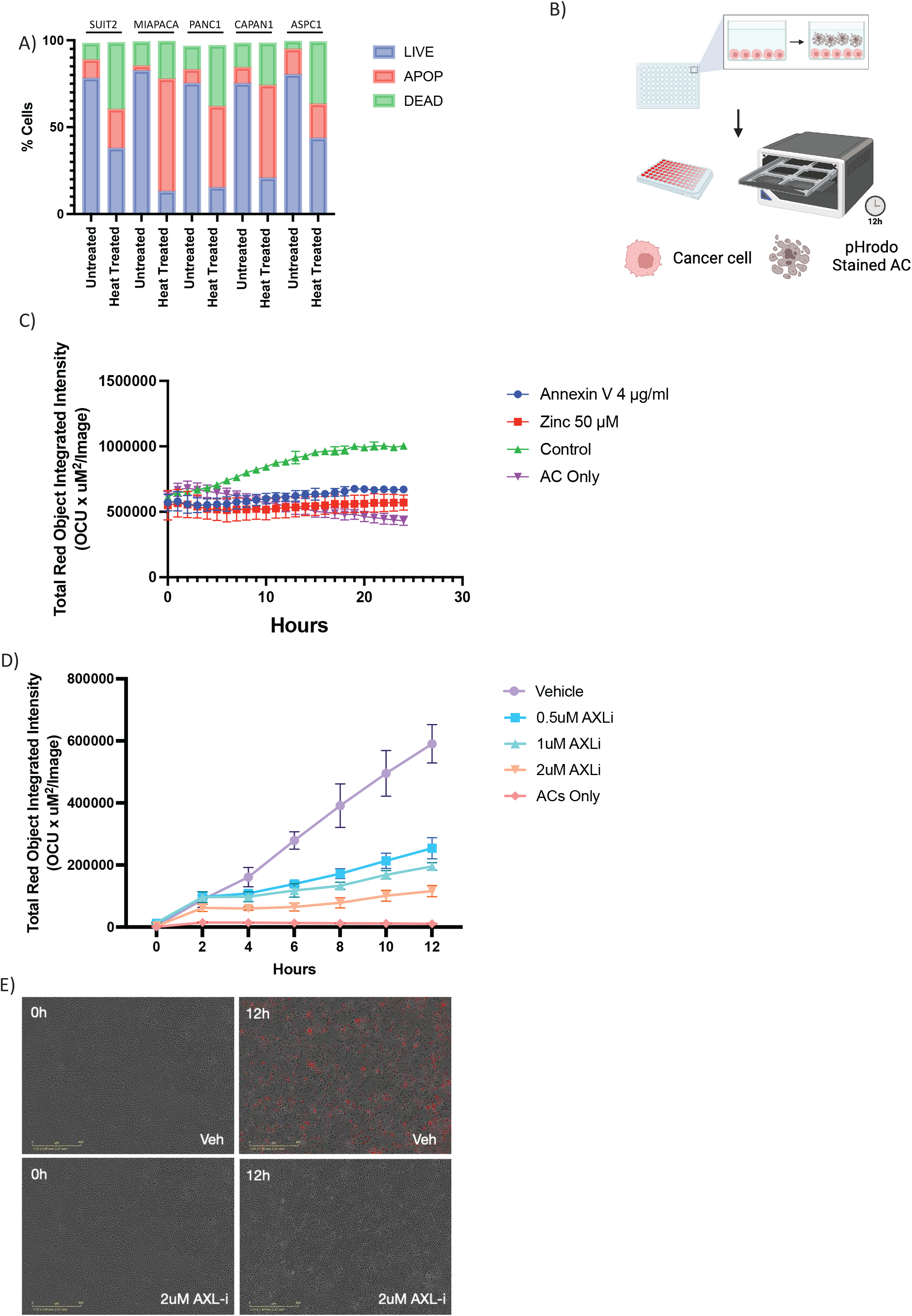
(A) Proportions of live, apoptotic, and dead cells before and after 30 minutes of 56°C heat treatment as measured by proprium iodide and annexin V with flow cytometry. (B) Schematic. SUIT2 cells were co-cultured with pHrodo labeled apoptotic cells, imaged at designated time points and measured for fluorescence activity. showing (C) Incucyte analysis of pHrodo activity in co-culture system treated with phosphatidylserine antibody Annexin V, efferocytosis inhibitor Zinc69391, or control over time. (D) Incucyte analysis of pHrodo activity in co-culture system treated with increasing dosages of AXL-i BGB324. (E) Representative images of (D).

**Supplementary Figure 2.**
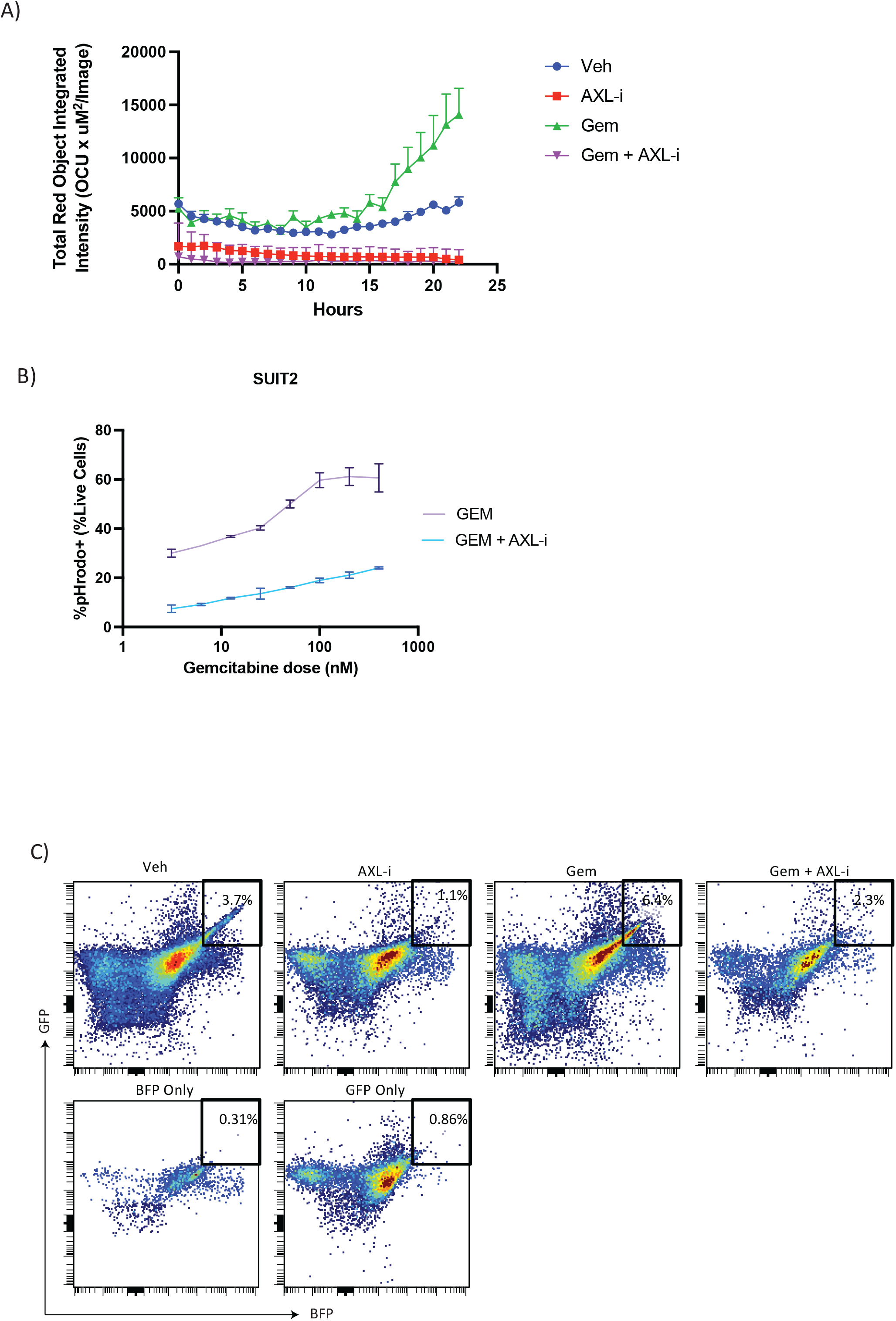
(A) Fluorescence activity over time as measured by Incucyte of pHrodo stained SUIT2 cells treated with AXL-i BGB324, gemcitabine, or both. (B) Percent pHrodo^+^ SUIT2 cells with increasing gemcitabine doses with or without AXL-i BGB324. (C) Representative flow plots and gating strategy of Figure 3E using fluorescence minus one controls.

**Supplementary Figure 3.**
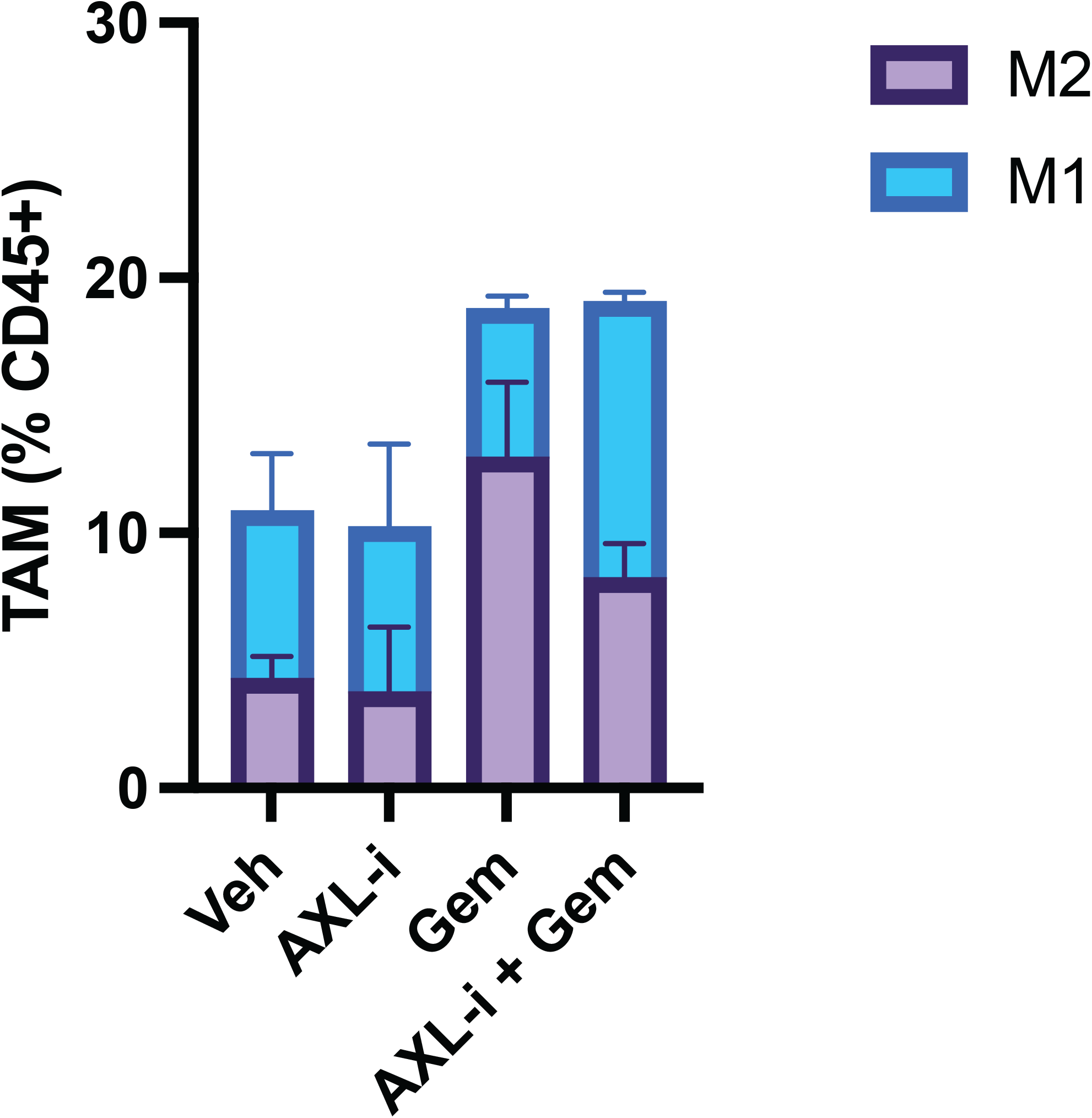
(A) Proportion of M1 and M2 macrophages by treatment group by single cell RNA analysis on treated KPC tumors from C57BL/6 mice.

